# MetaClaw: an auditable AI agent for end-to-end, multi-directional metagenomic and multi-omics analysis

**DOI:** 10.64898/2026.07.21.739769

**Authors:** Haohong Zhang, Zuqi Li, Philip Naderev Panuringan Lagniton, Zikang Wang, Linfeng Zhao, Wenxuan Li, Pengmin Duan, Xiaosen Jiang, Kang Ning

**Affiliations:** Key Laboratory of Molecular Biophysics of the Ministry of Education, Hubei Key Laboratory of Bioinformatics and Molecular-imaging, Center of AI Biology, Department of Bioinformatics and Systems Biology, College of Life Science and Technology, Huazhong University of Science and Technology, Wuhan 430074, Hubei, China; BGI Genomics, Shenzhen, China

## Abstract

End-to-end omics analysis requires more than selecting tools: a usable agent must bind data correctly, execute long workflows without blocking, preserve provenance and recover the biological conclusions that motivate an analysis. Existing LLM-driven bioinformatics agents automate parts of this process, but their operational dependencies and conclusion-level validity are often unclear. Here we present MetaClaw, an auditable agent that maps a user request to a registered workflow, executes standardized upstream processing on FlowHub, and runs study-specific downstream analyses in network-isolated OpenClaw containers. A YAML registry and an explicit plan–submit–poll–finalise lifecycle record file bindings, parameters, scripts, environments and outputs in per-job bundles. Across the full cohorts of four published studies (769 metagenomic profiles), MetaClaw recovered 4/4 sorghum marker groups, 3/3 RRMS features, 4/5 canonical CRC markers among the top 20 classifier features and 5/5 permafrost marker groups. In 45 model-by-prompt runs, upstream completion was consistent whereas downstream validity depended on the backend and instruction detail; three decoy-tested endpoints showed no significant differences. In 48 ablation sessions, removing the registry, planning loop or manifest caused distinct losses, with registry removal increasing time, tool calls and token cost. MetaClaw therefore connects standardized upstream execution, local analytical flexibility and conclusion-level validation in a rerunnable framework for metagenomic and microbiome multi-omics analysis.

## 1 Introduction

Across modern omics disciplines, converting raw measurements into biologically defensible conclusions requires long, multidirectional chains of interdependent analytical decisions. These decisions include read or feature quality control, host or contaminant removal, profiling, reference-database selection, statistical modelling, machine-learning evaluation, multi-omics integration and figure generation. Shotgun metagenomics is a demanding example: benchmarking studies have shown that analytical choices can substantially affect sensitivity, false-positive relative abundance, genome-reconstruction quality and functional coverage even when the input reads are held constant [1–3]. Workflow systems such as Snakemake [4] and metagenomic pipelines such as Sunbeam [5], MEDUSA [6] and UMGAP [7] reduce the engineering burden and improve computational reproducibility. However, once specified, these systems execute predefined graphs; they do not themselves define study-specific biological objectives, select or justify analyses, adapt downstream statistics to cohort structure, or assess whether a rerun reproduces the source study’s conclusions. Reproducibility also remains contingent on correct configuration of environments, parameters, reference databases and cross-tool hand-offs. Analysts must nevertheless choose appropriate workflows, connect tools and databases, adapt parameters to each cohort and translate computational results into biologically meaningful conclusions.

Large language model (LLM)-driven agents have recently begun to automate parts of this process. AutoBA showed that an autonomous agent could plan a multi-omics workflow from a high-level objective and recover from runtime failures through automated code repair [8]. BIA [9] and CellAgent [10] extended agentic analysis to single-cell RNA sequencing. BioAgents combines fine-tuning on tool documentation with retrieval-augmented generation to construct end-to-end workflows [11], whereas Biomni provides a general-purpose agentic environment containing more than 150 biomedical tools and databases and achieves strong zero-shot performance across benchmark tasks [12]. Despite this rapid progress, recent reviews consistently identify unstable multi-step reasoning, imperfect retrieval, weak biological grounding and limited reproducibility as major challenges [13, 14].

A central unresolved question is whether such agents can reproduce complete published studies rather than merely assemble plausible pipelines or execute code successfully. Existing evaluations typically assess step-level capabilities on curated tasks, such as BixBench [15], or whether an agent can construct an appropriate workflow, as in BioMaster [16]. Even when a system is described as end to end, validation often stops at successful execution, production of expected files or agreement on a single quantitative metric. These criteria do not establish whether an independent reanalysis recovers the principal biological conclusions of the original study.

This limitation is particularly evident in metagenomics and microbiome-centred multi-omics, where analyses combine heterogeneous data types and input formats, large raw sequencing datasets and privacy-sensitive biological information. Unlike many agent demonstrations that focus on single-cell or bulk-transcriptomic analyses, independent reproduction of published metagenomic and paired multi-omics studies remains rare. Such analyses require coordinated reasoning across taxonomic profiling, functional annotation, statistical inference, machine learning and cross-omics integration, making them a stringent test of whether an AI agent can faithfully reproduce biological findings. The missing component is therefore not simply another metagenomic pipeline, but an auditable (full execution traceable for peer review and privacy compliance) execution paradigm in which prompt-visible instructions, tool contracts, execution boundaries and generated outputs are transparent, inspectable and reusable.

Here we present MetaClaw, a bioinformatics AI agent organized around five linked contributions. First, it splits metagenomic analysis into a deterministic FlowHub upstream tool flow and a customizable OpenClaw downstream skill container. A single YAML pipeline registry couples the two layers. Second, per-job bundles archive FlowHub specifications, skill scripts, pinned Dockerfiles, parameters and outputs, while downstream containers run without network access. Third, four case-study reproductions spanning 769 metagenomic profiles test whether the framework recovers principal biological signals across sorghum drought, RRMS saliva, CRC stool and permafrost thaw. Fourth, 45 model-by-prompt runs and decoy perturbations test downstream reproducibility across changes in backend, instruction detail and irrelevant inputs. Fifth, 48 independent ablation sessions test how the registry, planning loop and per-job manifest contribute to completeness and resource use. Together, this design connects modular execution, security, rerunnable provenance, conclusion-level validation, perturbation robustness and operational efficiency.

## 2 Results

### 2.1 MetaClaw pipelines and skills

MetaClaw provides a modular, container-orchestrated framework that standardizes microbiome analysis while preserving study-specific flexibility (**Fig. 1**). The framework separates deterministic upstream processing from configurable downstream interpretation, thereby reducing ad hoc scripting while improving provenance tracking and execution reproducibility across shotgun metagenomic, amplicon and deep-learning-enabled analyses. Each analysis is assembled from a pipeline registry that couples a defined upstream FlowHub flow to a set of downstream OpenClaw skills within a Docker container. New analytical capabilities can therefore be added, or existing capabilities exchanged, without altering the language-model prompt itself.

**Figure 1:**
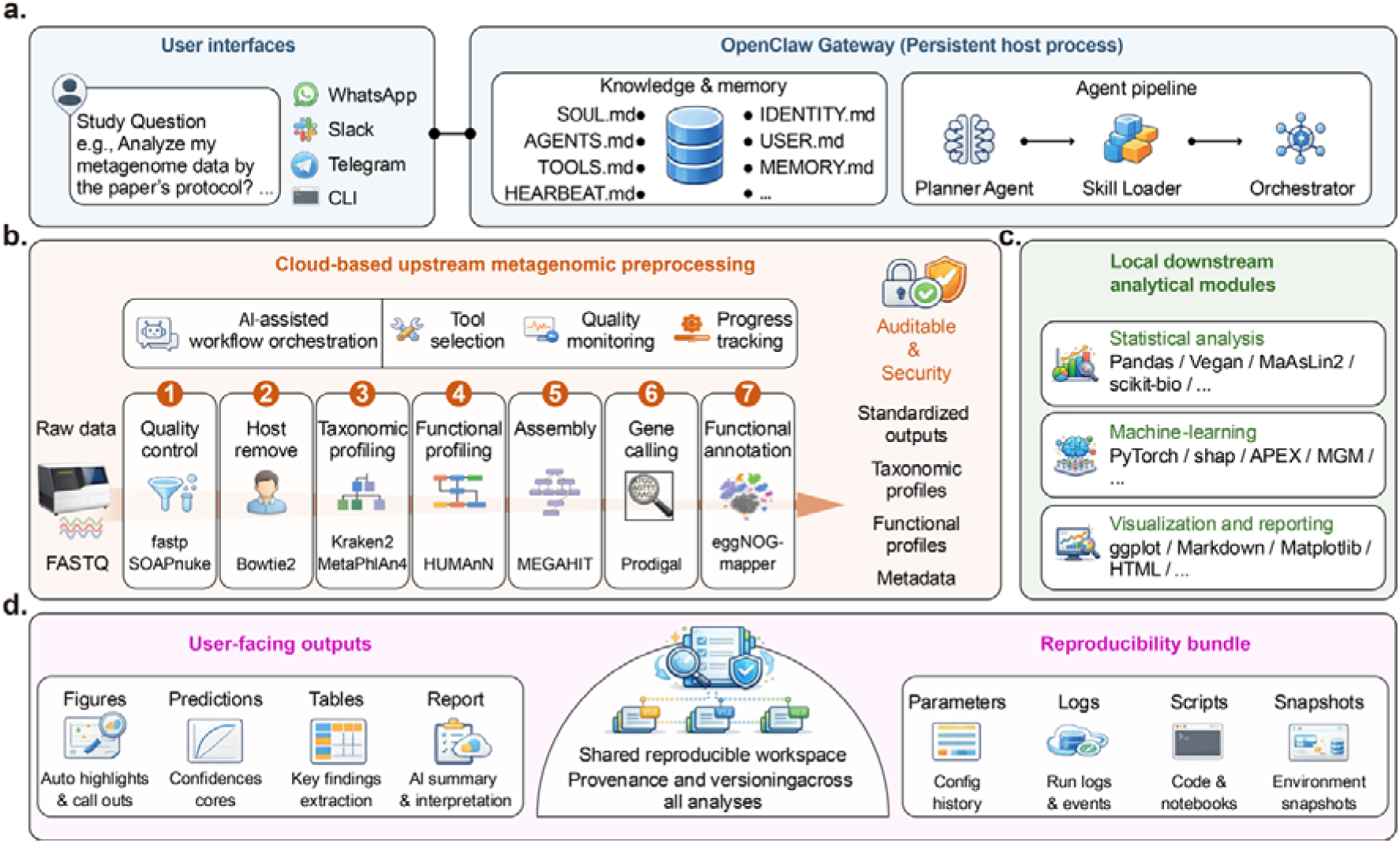
MetaClaw system architecture. A thin LLM-driven gateway connects a chat front-end to two heterogeneous execution back-ends through a single on-host job workspace. **a.** Gateway layer. A persistent OpenClaw process loads workspace files defining behavior (SOUL.md, AGENTS.md, TOOLS.md, HEARTBEAT.md, IDENTITY.md, USER.md, MEMORY.md) and exposes a Planner Agent, Skill Loader and Orchestrator; the gateway is reachable from common chat clients without code changes. **b.** Upstream back-end. QC, host-read decontamination, taxonomic and functional profiling, and assembly are delegated to versioned FlowHub flows invoked under a four-phase plan– submit–poll–finalize lifecycle; raw FASTQ reads stay on FlowHub. **c.** Downstream back-end. Statistical testing, machine-learning prediction, visualization and report generation run inside one of three pinned OpenClaw container images launched with *--network none* and *--cap-drop ALL*. **d.** Shared workspace and pipeline registry. A single YAML registry file is the only coupling point; the LLM never enters either back-end.

Users can access MetaClaw through the web deployment at https://www.bioline.cloud or a configured OpenClaw client. For end-to-end analysis, users provide a FlowHub data path; MetaClaw selects the registered pipeline, confirms input bindings, monitors upstream execution and stages completed outputs for downstream analysis (**Methods 5.2**).

MetaClaw currently offers five pipeline configurations for different analytical scenarios (**Table 1**; **Supplementary Table 1**). A pipeline may couple one FlowHub flow (upstream) to a skill container (downstream) through the pipeline registry, or focus solely on upstream or downstream analysis. Metagenomics-full is the default pipeline, proceeding from raw paired-end reads to taxonomic profiling, downstream statistics, diversity and differential-abundance testing, visualization and reporting. Upstream-only exposes the same cloud-based preprocessing for users who prefer external downstream analysis, whereas downstream-only reanalyzes precomputed abundance or pathway tables using the same analytical framework. Amplicon-ASV provides a complete 16S/ITS workflow from denoising to taxonomic assignment and functional prediction. Downstream-DL adds deep-learning modules for disease classification, profile generation, antimicrobial peptide prediction, biosynthetic gene cluster detection and antibiotic-resistance annotation. Together, these pipelines allow MetaClaw to move from raw sequence data to downstream interpretation within one reproducible framework.

**Table 1:** Overview of all available pipelines in MetaClaw. It lists the analytical scope of each pipeline, the required starting data, and the principal deliverables generated at each stage, spanning various upstream and downstream analysis combinations. More details of the pipelines can be found in Supplementary Table 1.

| Pipeline | Description | Input type | Main outputs |
| --- | --- | --- | --- |
| Metagenomics-full | Full shotgun metagenomics | Raw paired-end FASTQ | Publication-ready statistical reports and curated abundance tables |
| Upstream-only | Standardized upstream processing only | Raw FASTQ + metadata | Taxonomic profiles plus QC and host-removal reports for external analysis |
| Downstream-only | Reanalysis of preprocessed data | Abundance tables, pathway profiles, or sequence data | Statistical summaries, figures, and reports |
| Amplicon-ASV | 16S/ITS amplicon analysis | Amplicon sequencing reads | ASV tables, taxonomy, predicted function, downstream-ready matrices |
| Downstream-DL | Deep-learning-enabled downstream analysis | Microbiome profiles and sequence-based inputs | Predictions, annotations, and model-derived outputs |

The upstream stage of each pipeline runs as one of seven versioned FlowHub flows invoked through the *fkit* interface (**Supplementary Table 2**). These flows cover 1) host removal and MetaPhlAn 4 profiling, 2) contig assembly, 3) genome-resolved binning, 4) protein annotation against public databases, 5) antibiotic-resistance screening, 6) organelle assembly and 7) read subsampling. Downstream analytical capability is organized as a catalogue of skills, each pairing a SKILL.md contract with a customized reference script (**Supplementary Table 3**). The catalogue spans statistics, microbiome ecology, metabolomics, machine learning, figure generation and reporting, and includes the deep-learning skills MGM, APEX, BGC-Prophet and ONN4ARG [17–20]. This modular design allows capabilities to be added by extending individual workflows or skills without modifying the overall execution framework.

### 2.2 End-to-end reproduction of heterogeneous metagenomic studies

We next evaluated whether MetaClaw could reproduce complete metagenomic studies rather than isolated analytical tasks. The unified workflow connects input preprocessing, read-based taxonomic profiling, downstream microbiome analysis, study-specific validation and manuscript-ready outputs (**Fig. 2**). It accepts raw sequencing reads or verified preprocessed taxonomic profiles, performs quality control and host-read removal when required, and generates MetaPhlAn 4 species or species-level genome bin (SGB) profiles. Individual sample profiles are then merged into harmonized abundance matrices, providing a common analytical representation across datasets with different sample types, designs and biological contexts.

**Figure 2:**
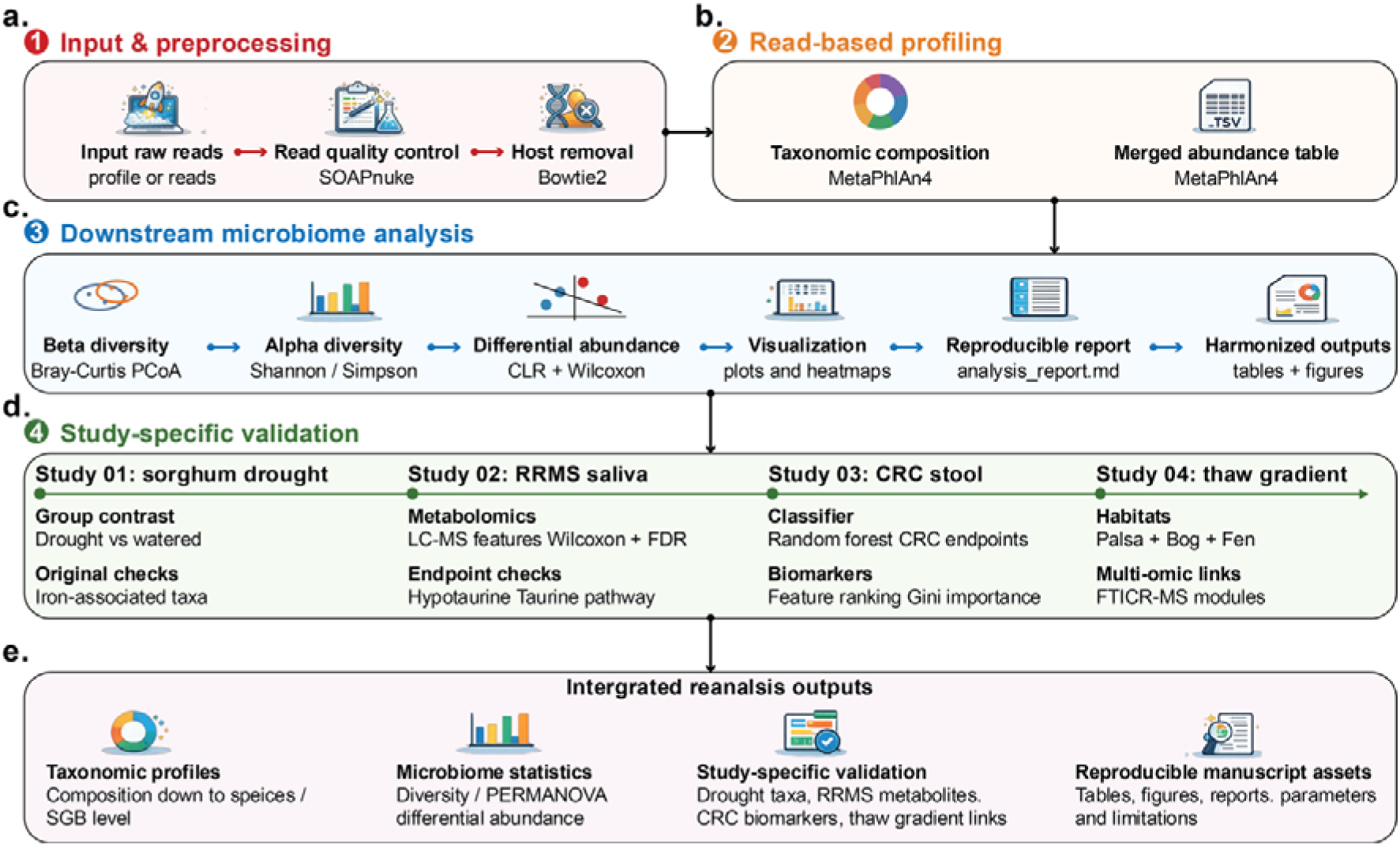
Unified agent workflow for end-to-end reproduction of metagenomic studies. **a.** Input and preprocessing module. The workflow accepts either raw sequencing reads or verified preprocessed taxonomic profiles, followed by read quality control using SOAPnuke and host-read removal using Bowtie2 when required. **b.** Read-based profiling module. Quality-controlled microbial reads are profiled using MetaPhlAn 4 to generate taxonomic composition profiles, which are merged into unified abundance tables for downstream analysis. **c.** Shared downstream microbiome analysis module. The merged abundance tables are analyzed through alpha-diversity testing, beta-diversity ordination, CLR-based differential abundance analysis, visualization, reproducible report generation, and harmonized output formatting. **d.** Study-specific validation module. The shared workflow branches into benchmark-specific checks: drought and compartment contrasts with publication-anchored taxa for the sorghum drought study; LC-MS metabolite analysis and hypotaurine/taurine endpoint checks for the RRMS saliva multi-omics study; random-forest classification and biomarker ranking for the CRC stool metagenome study; and habitat-gradient diversity, FTICR-MS metabolite modules and greenhouse-gas-associated marker taxa for the permafrost thaw-gradient study. **e.** Integrated reanalysis outputs. Final outputs include species- or SGB-level taxonomic profiles, microbiome statistics, study-specific validation results and reproducible manuscript assets, including tables, figures, reports and parameters. CLR, centered log-ratio; LC–MS, liquid chromatography– mass spectrometry; RRMS, relapsing-remitting multiple sclerosis; CRC, colorectal cancer;

To combine analytical consistency with study-specific evaluation, the downstream stage comprises a shared analytical layer and study-specific validation modules. The shared layer performs alpha-diversity analysis, beta-diversity ordination, centred log-ratio (CLR)-based differential-abundance testing, visualization, harmonised output generation and reproducible reporting (**Fig. 2c**). Study-specific modules then evaluate publication-anchored endpoints: drought- and compartment-associated taxa in the sorghum benchmark; LC-MS metabolite features and hypotaurine or taurine endpoints in the RRMS saliva benchmark; random-forest classification and canonical marker recovery in the CRC benchmark; and habitat-associated microbiome–metabolite structure in the permafrost benchmark. This organization separates general microbiome analysis from benchmark-specific validation, a key protocol-like feature of the workflow.

#### 2.2.1 Reproduction of a sorghum rhizosphere drought-stress metagenomic study

We first tested whether MetaClaw could reproduce an environmental metagenomic study [21] in which community structure was shaped by both plant compartment and drought treatment. The complete benchmark contained 47 sorghum-associated metagenomes: 11 drought-rhizosphere, 12 watered-rhizosphere, 12 drought-bulk-soil and 12 watered-bulk-soil samples. Starting from sequencing reads, the agent completed quality control, sorghum host-read removal, MetaPhlAn 4 taxonomic profiling, merged-table construction and downstream ecological analysis (**Fig. 3a**).

**Figure 3:**
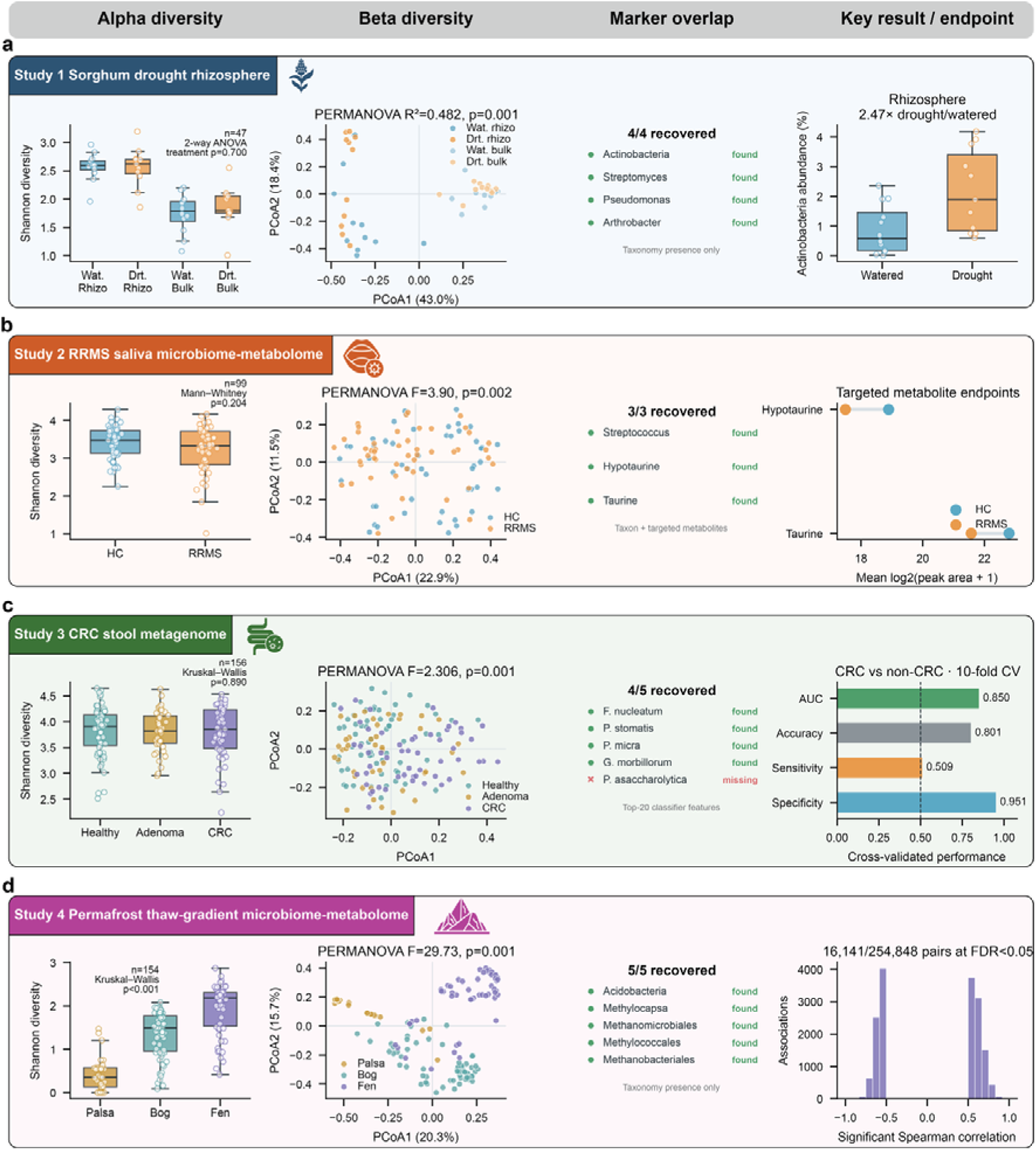
MetaClaw reproduction of four microbiome and multi-omics benchmark studies using the full benchmark datasets. Within each study row, panels show alpha diversity, beta diversity, publication-anchored feature recovery and a study-specific endpoint. **a.** Sorghum drought-rhizosphere metagenomics (n = 47). The four-level treatment-compartment PERMANOVA model explained 48.2% of community variation (F = 13.34, p = 0.001). All four anchored marker groups were detected in the complete species table. Mean rhizosphere Actinobacteria abundance was 2.180% in drought and 0.884% in watered samples, a 2.47-fold difference. **b.** RRMS saliva microbiome-metabolome profiling used 99 metagenomes (50 healthy controls and 49 RRMS) and 20 metabolomics samples. Shannon diversity did not differ significantly (Mann-Whitney p = 0.204), whereas community composition differed by PERMANOVA (F = 3.90, p = 0.002). All three anchored features were recovered. Hypotaurine decreased in RRMS (RRMS-HC log2FC = -1.402; two-sided p = 0.0939; one-sided p = 0.0469), and taurine also decreased (log2FC = -1.227; two-sided p = 0.00357). **c.** CRC stool metagenomics comprised 156 samples (61 Healthy, 42 Adenoma and 53 CRC). Shannon diversity did not differ among groups (Kruskal-Wallis p = 0.890), whereas beta diversity differed by PERMANOVA (F = 2.306, p = 0.001). Four of five canonical CRC taxa ranked among the top 20 classifier features, and the CRC-versus-non-CRC random forest achieved AUC = 0.850, accuracy = 0.801, sensitivity = 0.509 and specificity = 0.951 in 10-fold cross-validation**. d.** Permafrost profiling produced 467 species profiles; habitat-aware analyses used 154 labelled profiles (33 Palsa, 63 Bog and 58 Fen). Mean Shannon diversity increased from Palsa (0.415) to Bog (1.344) and Fen (1.936), and habitat-associated composition differed by PERMANOVA (F = 29.73, p = 0.001). All five anchored clade names occurred in the complete taxonomy table; these are taxonomy-presence observations, not functional or gas-flux measurements. Of 254,848 tested microbe-metabolite correlations in the paired analysis, 16,141 passed FDR < 0.05. Feature-recovery definitions were fixed to the corresponding data type: presence in the complete taxonomy table for Studies 1 and 4, directional recovery for Study 2, and top-20 classifier ranking for Study 3. Wat., watered; Drt., drought; Rhizo, rhizosphere; HC, healthy control; RRMS, relapsing-remitting multiple sclerosis; CRC, colorectal cancer; AUC, area under the receiver operating characteristic curve; PCoA, principal coordinates analysis; PERMANOVA, permutational multivariate analysis of variance; FDR, false discovery rate.

In the complete cohort, drought treatment did not produce a broad alpha-diversity shift (two-way ANOVA treatment effect, p = 0.70), whereas rhizosphere samples remained more diverse than bulk-soil samples. Bray-Curtis ordination separated the four treatment-compartment groups, whose PERMANOVA model explained 48.2% of community variation (R2 = 0.482, F = 13.34, p = 0.001; beta-diversity panel of **Fig. 3a**). Group dispersions were heterogeneous (betadisper p = 0.0198), so the PERMANOVA result reflects a combination of compositional location and dispersion differences.

All four publication-anchored marker groups—Actinobacteria, Streptomyces, Pseudomonas and Arthrobacter—were present in the complete species table (feature-recovery panel of **Fig. 3a**). Differential analysis identified Streptomyces capitiformicae and Glycomyces albidus as drought-enriched rhizosphere species at FDR = 0.058. Mean rhizosphere Actinobacteria abundance was 2.180% under drought and 0.884% under watered conditions, corresponding to a 2.47-fold enrichment (key-endpoint panel of **Fig. 3a**). Detection of a marker name denotes taxonomic recovery and does not establish iron-metabolism function.

Overall, the full sorghum benchmark recovered 4/4 publication-anchored marker groups, a 2.47-fold drought enrichment of rhizosphere Actinobacteria and strong treatment-compartment community structure (PERMANOVA R2 = 0.482, p = 0.001). The differential signal was concentrated in Actinobacteria, with two drought-enriched species at FDR < 0.10 and no significant drought-associated species in bulk soil.

#### 2.2.2 Reproduction of an RRMS saliva microbiome–metabolome study

We next evaluated whether MetaClaw could reproduce a disease-associated oral microbiome-metabolome study [22] using the complete set of 99 saliva metagenomes from 50 healthy controls and 49 patients with relapsing-remitting multiple sclerosis (RRMS) and 20 LC-MS metabolomics samples. MetaClaw executed the study-specific workflow, including read quality control, human host-read removal, MetaPhlAn 4 species-level taxonomic profiling, construction of merged microbial abundance tables, microbial diversity analysis, species-level differential-abundance testing and metabolite-level analysis (**Fig. 3b**).

In the full 99-sample metagenomic cohort, Shannon diversity did not differ significantly between healthy controls and RRMS samples (Mann-Whitney p = 0.204; alpha-diversity panel of **Fig. 3b**). Simpson diversity was marginally lower in RRMS (p = 0.047), although this nominal result was one of four alpha-diversity metrics tested. By contrast, Bray-Curtis community composition differed between groups by PERMANOVA (F = 3.90, p = 0.002; beta-diversity panel of **Fig. 3b**), indicating a community-level disease association without a broad richness or Shannon-diversity shift.

Publication-anchored validation recovered all three predefined features: Streptococcus, hypotaurine and taurine (feature-recovery panel of **Fig. 3b**). Streptococcus salivarius and Streptococcus sanguinis were depleted in RRMS at FDR = 0.00350 and 0.0447, respectively. In the metabolomics analysis, hypotaurine was lower in RRMS (RRMS-HC log2FC = -1.402; HC n = 8, RRMS n = 7; two-sided p = 0.0939; prespecified one-sided p = 0.0469), while taurine showed a clearer decrease (log2FC = -1.227; HC n = 9, RRMS n = 9; two-sided p = 0.00357; key-endpoint panel of **Fig. 3b**). No metabolite passed FDR < 0.05 across the untargeted 964-metabolite screen.

Overall, the RRMS benchmark recovered 3/3 publication-anchored features and significant community separation in the complete metagenomic cohort (PERMANOVA p = 0.002). The targeted metabolite results were directionally concordant, with hypotaurine supported by the prespecified one-sided test.

#### 2.2.3 Reproduction of a colorectal cancer stool metagenome biomarker study

We next tested whether MetaClaw could reproduce a stool metagenome-based disease-classification study [23] of colorectal cancer (CRC) using the complete 156-sample French cohort: 61 healthy controls, 42 adenoma samples and 53 CRC samples. The agent analyzed preprocessed, host-depleted metagenomic reads and performed MetaPhlAn 4 species-level profiling, merged abundance-table construction, diversity analysis, canonical marker ranking and random-forest classification of CRC status (**Fig. 3c**).

Alpha diversity did not differ among Healthy, Adenoma and CRC samples in the exported full-cohort table (Kruskal-Wallis p = 0.890; alpha-diversity panel of **Fig. 3c**), whereas beta diversity differed by PERMANOVA (F = 2.306, p = 0.001; beta-diversity panel of **Fig. 3c**). Four of five canonical CRC-associated taxa ranked among the top 20 random-forest features: Peptostreptococcus stomatis (#1), Fusobacterium nucleatum (#2), Parvimonas micra (#4) and Gemella morbillorum (#5). Porphyromonas asaccharolytica was not recovered among the profiled classifier features (feature-recovery panel of **Fig. 3c**).

The full-cohort random forest distinguished CRC from non-CRC with an AUC of 0.850 in stratified 10-fold cross-validation (key-endpoint panel of **Fig. 3c**). Accuracy was 0.801, sensitivity was 0.509 and specificity was 0.951. The lower sensitivity than specificity indicates that the model was better at ruling out CRC than detecting all CRC cases.

#### 2.2.4 Reproduction of a permafrost thaw-gradient microbiome-metabolome study

We next evaluated whether MetaClaw could reproduce an environmental multi-omics study of microbiome-metabolite structure across a permafrost thaw gradient [24]. The complete taxonomic output contained 467 peat metagenomic profiles. Habitat-aware analyses used the 154 profiles with Palsa, Bog or Fen labels (33, 63 and 58 samples, respectively), and paired FTICR-MS data were used for multi-omics integration. The agent performed MetaPhlAn 4 profiling, microbial diversity analysis, taxonomy-based marker screening, metabolite feature testing and microbe-metabolite correlation analysis (**Fig. 3d**).

Among the 154 habitat-labelled profiles, mean Shannon diversity increased from Palsa (0.415, n = 33) to Bog (1.344, n = 63) and Fen (1.936, n = 58; alpha-diversity panel of **Fig. 3d**). Bray-Curtis PCoA showed habitat-associated community structure, and PERMANOVA detected group differences (F = 29.73, p = 0.001; beta-diversity panel of **Fig. 3d**).

All five publication-anchored clade names—Acidobacteria, Methylocapsa, Methanomicrobiales, Methylococcales and Methanobacteriales—were present in the complete 467-profile taxonomy table (feature-recovery panel of Fig. 3d). The targeted candidate export itself contained three unique Acidobacteria taxa across three habitats and no entries from the other four anchor groups, illustrating that taxonomy-table detection and candidate-screen output are distinct criteria. Neither criterion demonstrates methane metabolism, gene expression or measured CO2/CH4 fluxes.

The full metabolite output contained 5,792 unique FTICR-MS features represented in 16,258 pairwise habitat-comparison rows. Of these, 3,868 comparison rows, corresponding to 2,721 unique features, passed FDR < 0.05: 1,237 rows for Bog versus Palsa, 676 for Fen versus Palsa and 1,955 for Fen versus Bog. These feature-level results provide association evidence rather than direct functional or flux confirmation.

The paired multi-omics output contained 254,848 microbe-metabolite Spearman correlations, of which 16,141 (6.33%) passed FDR < 0.05 (key-endpoint panel of **Fig. 3d**). The strongest absolute correlation in the exported table was 0.987. These associations remain correlative and do not identify causal microbial-metabolite mechanisms.

### 2.3 Robustness across models, prompts and noisy input perturbations

To investigate whether MetaClaw’s downstream conclusions remain valid when the LLM backend, prompt specification or irrelevant input context changes, we performed a 45-run 3 × 3 model-by-prompt benchmark together with decoy perturbations. Across the factorial benchmark, all three LLM models — Qwen (qwen3.7-max), DeepSeek (deepseek-v4-pro) and Doubao (doubao-seed-2-0-pro-260215)—completed the deterministic upstream pipeline, but downstream validity diverged by model and prompt granularity (Fig. 4a). Profile merging remained nearly universal. Doubao’s alpha- and beta-diversity outputs were invalid when diversity was computed across taxa rather than samples. Qwen was the most robust overall: it achieved the highest valid completion rates across prompt conditions and was the only model to recover the requested multifactor PERMANOVA model. DeepSeek responded nonlinearly to prompt granularity, performing best with the full prompt and worst with the medium prompt. We compared DeepSeek and Qwen within the Full, Medium and Simple conditions using pooled raw replicate-level values and Mann–Whitney U tests on shared valid samples and features, excluding zero-Shannon alpha-diversity rows before pooling (Fig. 4b). The only significant contrast was beta diversity in the Simple condition (p = 0.0055). All other panel B comparisons were not significant: alpha diversity in Full, Medium and Simple (p = 0.9489, 0.1964 and 0.0775), beta diversity in Full and Medium (p = 1.0000 and 0.1113), and species log2FC in Full, Medium and Simple (p = 0.6605, 0.7503 and 0.4848).

**Figure 4:**
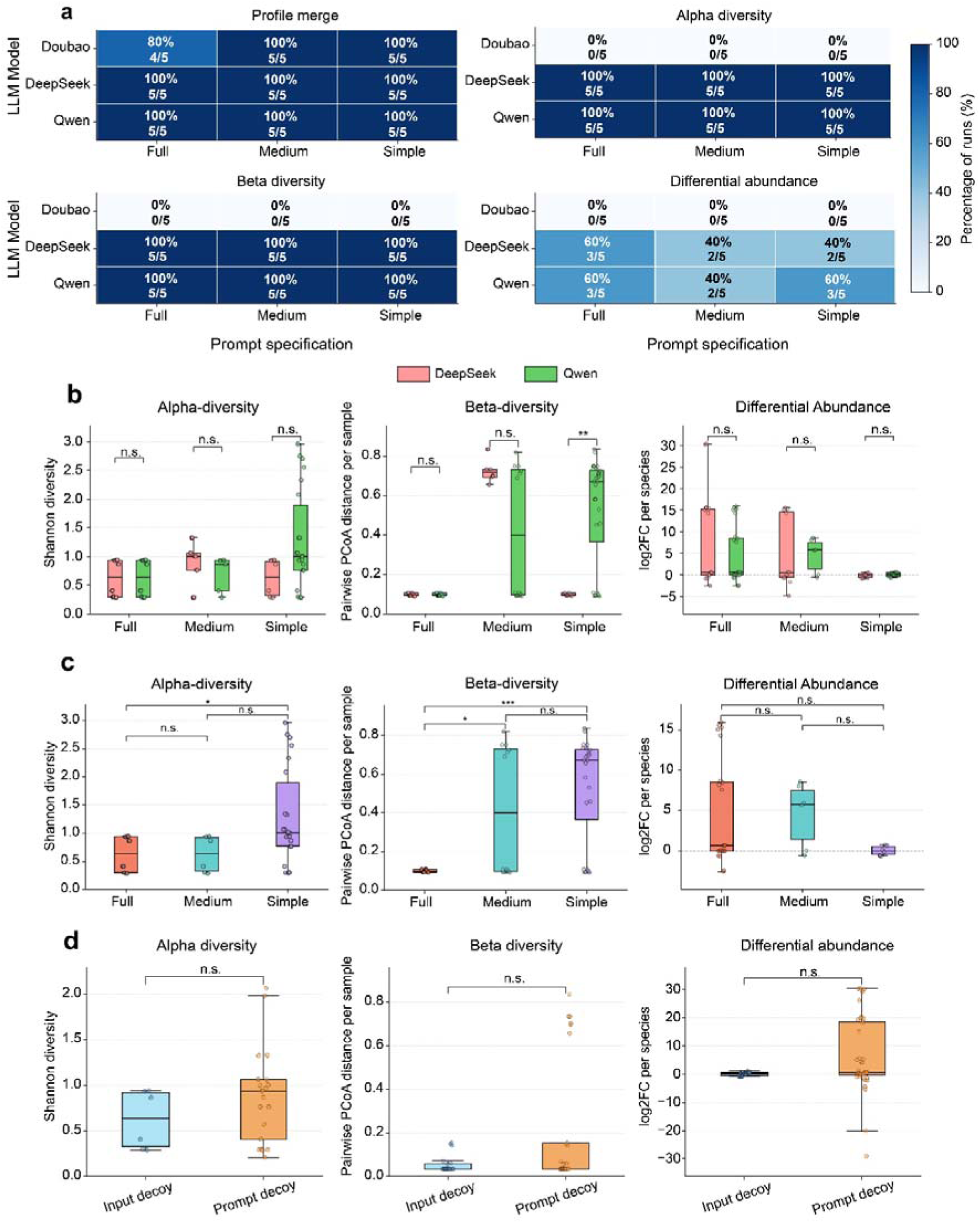
Composite robustness figure showing valid completion across model and prompt perturbations, replicate-level consistency for the valid downstream analyses, and study-selection fidelity under mixed-folder decoy conditions. a. Completion-rate heatmaps across pipeline stages. b. Pooled raw-value comparisons of DeepSeek and Qwen across Full, Medium, and Simple using Mann –Whitney U tests on shared valid samples and features. c. Comparison between prompt granularity using Qwen. d. Pooled raw-value comparisons of Qwen input-decoy and prompt-decoy conditions using shared valid samples for alpha and beta, and shared species/features for log2FC, with Mann– Whitney U tests.

Within Qwen, differential-abundance estimates were robust to prompt granularity, whereas diversity summaries were more sensitive (Fig. 4c). Shannon alpha diversity did not differ between Full and Medium (raw p = 0.9792; Holm p = 0.9792) or Medium and Simple (raw p = 0.0775; Holm p = 0.1551), but differed between Full and Simple (raw p = 0.0053; Holm p = 0.0160). Beta diversity differed between Full and Medium (raw p = 0.0226; Holm p = 0.0452) and between Full and Simple (raw p = 3.87 × 10^-5; Holm p = 1.16 × 10^-4), whereas Medium and Simple did not differ (raw p = 0.6503; Holm p = 0.6503). Species log2FC did not differ for any prompt pair after Holm correction (Full versus Medium, Holm p = 0.8151; Medium versus Simple, Holm p = 0.3961; Full versus Simple, Holm p = 0.4060). Qwen was therefore most stable for differential abundance and showed prompt sensitivity mainly in alpha- and beta-diversity summaries, especially in contrasts involving Full and Simple. Because the benchmark used a 10% read subsample and the Simple prompt was the least prescriptive condition, the remaining differences probably reflect a combination of prompt-granularity effects and sampling variability. The shorter prompt left more procedural detail implicit, allowing stochastic variation in the sampled reads to propagate more readily into diversity summaries than under the more explicit prompts.

The decoy benchmark showed that the workflow was robust to irrelevant input and prompt perturbations. We applied the same shared-valid raw-value logic to Qwen input-decoy and prompt-decoy conditions **(Fig. 4d)**. We pooled raw replicate-level values across shared valid samples for alpha and beta diversity and across shared species or features for log2FC, excluding zero-Shannon alpha-diversity rows before pooling. Shannon alpha diversity and pairwise PCoA distance per sample did not differ between the decoy conditions (p = 0.2550 and 0.1770, respectively), and species log2FC was also not significant (p = 0.0712). Complete comparison statistics are provided in **Supplementary Table S4.**

### 2.4 Ablation tests of component contributions

To determine which scaffold components are necessary for valid and efficient execution, we compared the intact agent with variants lacking the planning loop, pipeline registry or per-job manifest (Fig. 5; Table 2). Completeness was evaluated using a weighted checklist score. Mean overall completeness was 0.85 ± 0.03 in the baseline, statistically indistinguishable in −Manifest (0.85 ± 0.04) and −Plan (0.84 ± 0.03), and lowest in −Registry (0.72 ± 0.08). The registry contrast did not reach significance on the trimmed ranks (p = 0.21, Mann–Whitney U, two-sided) because the arm was strongly bimodal — some runs reconstructed the pipeline fully while others failed outright, so between-replicate variance remained high. Reliability told the same story more sharply: across all twelve untrimmed replicates the baseline never fell below 0.72, whereas the registry ablation produced catastrophic, near-empty outputs in a third of its runs. At the category level the registry ablation lowered the MetaPhlAn-table, differential-abundance and figure scores, consistent with several replicates never reconstructing the upstream–downstream contract, while the narrative report was recovered in every arm.

**Figure 5:**
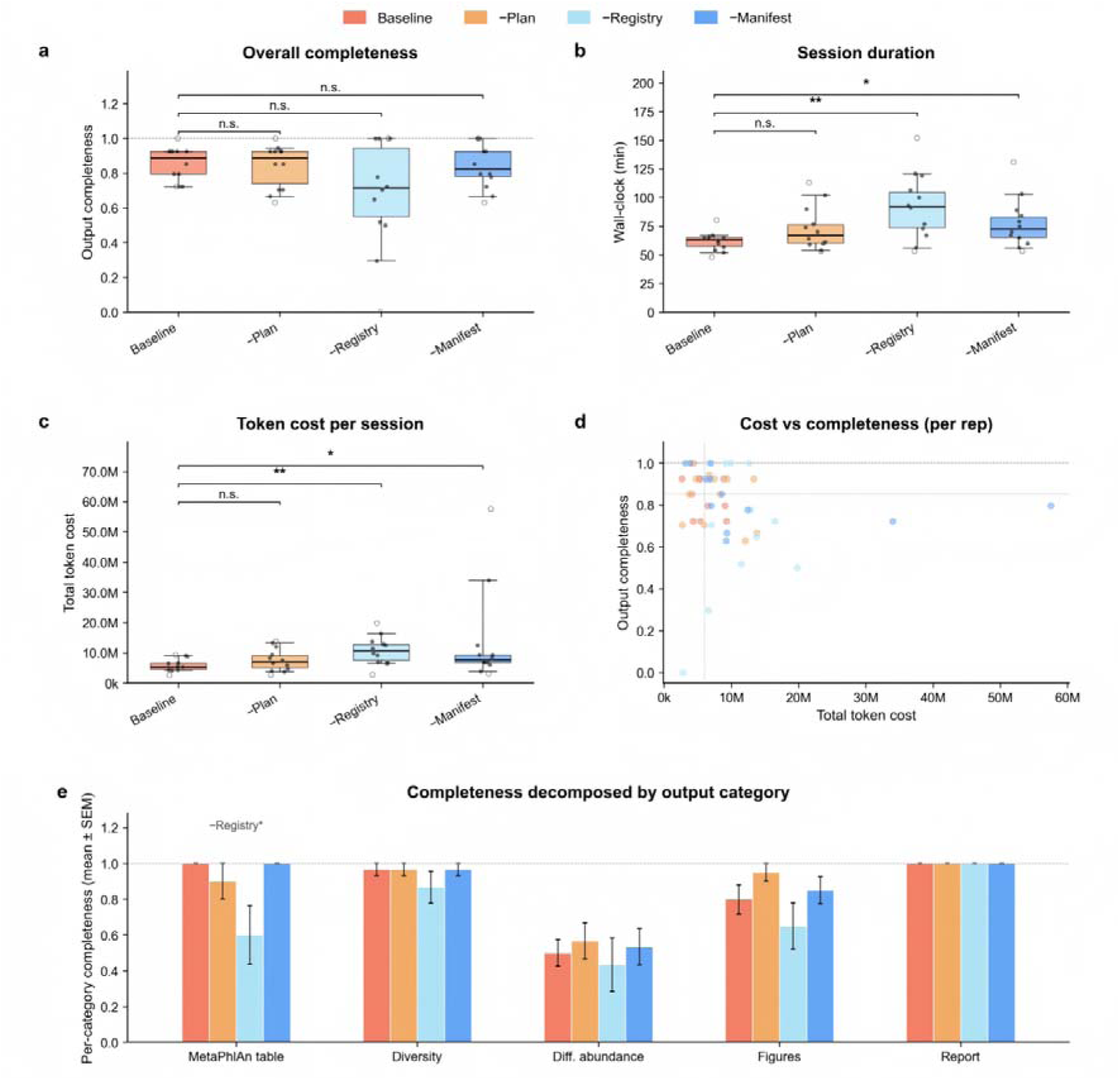
Effect of removing individual scaffold components from MetaClaw on the sorghum drought benchmark (n = 12 replicates per arm, trimmed to n = 10 by removing the single smallest and single largest values per arm and metric). **a.** Overall completeness per arm, shown as boxplots (median line + IQR box + min-max whiskers); solid dots are the kept replicates and open circles are the min/max trimmed out. **b.** Wall-clock session duration. **c.** Total token cost per session on a linear axis. **d.** Per-replicate overall completeness plotted against total token cost, with dotted lines marking the baseline mean on each axis. **e.** Completeness decomposed into five output categories (mean ± SEM on the trimmed values). Pairwise brackets in **a–c** show two-sided Mann– Whitney U p-values against baseline; *p < 0.05, **p < 0.01, ***p < 0.001, n.s. otherwise.

**Table 2:** Per-arm summary of the ablation experiment. Values are mean ± standard error of the mean on the trimmed n = 10 replicates per arm (single smallest and single largest of 12 replicates dropped). Significance markers reflect two-sided Mann–Whitney U p-values against baseline; *p < 0.05, **p < 0.01; n.s., not significant.

| Arm | Completeness | Walltime (min) | Tool calls | Total tokens (M) |
| --- | --- | --- | --- | --- |
| Baseline | $0.85 \pm 0.03$ | $61 \pm 2$ | $67 \pm 4$ | $5.99 \pm 0.56$ |
| $\text{-Plan}$ | $0.84 \pm 0.03$ (n.s.) | $71 \pm 5$ (n.s.) | $73 \pm 7$ (n.s.) | $7.56 \pm 1.03$ (n.s.) |
| $\text{-Registry}$ | $0.72 \pm 0.08$ (n.s.) | $90 \pm 7$ (**) | $95 \pm 6$ (**) | $10.65 \pm 1.06$ (**) |
| $\text{-Manifest}$ | $0.85 \pm 0.04$ (n.s.) | $75 \pm 5$ (*) | $69 \pm 6$ (n.s.) | $10.43 \pm 2.72$ (*) |

Session cost separated the arms more clearly than completeness did (Fig. 5b–c; Table 2). Mean wall-clock time increased from 61 ± 2 min at baseline to 90 ± 7 min in −Registry (p = 0.002) and 75 ± 5 min in −Manifest (p = 0.017), while −Plan was statistically indistinguishable from baseline. Tool invocations rose from 67 ± 4 in the baseline to 95 ± 6 in −Registry (p = 0.001) but not in the other arms. The cost in language-model tokens showed the same separation: mean total token cost was 5.99 ± 0.56 M in the baseline versus 10.65 ± 1.06 M in −Registry (p = 0.001) and 10.43 ± 2.72 M in−Manifest (p = 0.038), whereas −Plan (7.56 ± 1.03 M) did not differ significantly. Baseline replicates cluster in the low-cost, high-completeness corner, whereas −Registry replicates scatter along both axes, reflecting the arm’s combination of failed reconstructions and heavy exploratory token spend (Fig. 5d). Thus, the registry and manifest ablations move the cost–quality trade-off in different directions, i.e., registry loss raises cost while pushing a subset of runs to low completeness, whereas manifest loss preserves completeness but at a substantially higher and more variable token spend.

Disk use provided an additional operational signal that separated the registry arm from the others. Every −Registry replicate produced a job folder substantially larger than the approximately 5 MB norm observed in the other arms, ranging from a few hundred megabytes to tens of gigabytes. Inspection showed that the agent had downloaded raw upstream FlowHub outputs and, in several runs, software-environment caches that the normal pipeline contract would have retained on the cloud side. Two −Manifest replicates showed the same pattern, with folder sizes of 8.1 and 26 GB. The remaining−Manifest replicates and all baseline and −Plan replicates remained near 5 MB. Although these enlarged job directories still scored at or near baseline for completeness, their disk footprint would be prohibitive on metered storage.

Together, the ablations show that each scaffold layer contributes a different operational function. The registry encodes the upstream–downstream contract and prevents both exploratory reconstruction of pipeline settings and indiscriminate file retrieval. Its removal produced the lowest and most variable completeness, significantly increased wall-clock time, tool invocations and token cost, and inflated disk footprint by three to four orders of magnitude. The manifest carries the analytical hand-off between stages, and removing it significantly inflated token cost and session duration without measurably hurting the deliverable count, because the agent absorbs the missing index by re-reading the staging directory. The planning loop is the cheapest single layer to keep, producing the smallest change across every metric while still guarding against fragile runs.

## 3 Discussion

This study establishes five linked properties of MetaClaw. First, a single YAML registry couples deterministic FlowHub upstream processing to customizable OpenClaw downstream skills. Second, network-isolated downstream execution and per-job archival make the analytical procedure secure, inspectable and rerunnable. Third, four heterogeneous case studies test conclusion-level biological reproduction rather than workflow execution alone. Fourth, model, prompt and decoy perturbations quantify downstream robustness. Fifth, component ablations identify how the registry, planning loop and manifest affect completeness and resource use. Together, these tests distinguish architecture, security and reproducibility, biological reproduction, perturbation robustness and scaffold necessity as complementary properties of an agentic scientific system.

MetaClaw’s main architectural contribution is the division of an omics analysis into a deterministic upstream tool flow and a customizable downstream skill container, joined by a single YAML pipeline registry. Versioned FlowHub flows standardise quality control, host-read removal and profiling, whereas downstream skills provide local Python and R analyses for study-specific statistics, visualization and reporting. Because the registry is the only coupling point between the two domains, workflow selection, expected inputs and outputs, and container images are explicit rather than inferred at runtime.

Security and reproducibility are enforced at the job level rather than described after the analysis. Each job bundle archives the FlowHub specification, OpenClaw skill scripts, pinned Dockerfiles, execution manifest and outputs, making both the agent-visible instructions and the computational environment inspectable. Downstream containers run without network access and with Linux capabilities removed, while the LLM remains outside both computational backends. This design constrains common prompt-injection routes into the runtime and preserves the evidence needed to reconstruct what was planned, executed and produced.

The case analyses tested conclusion-level reproduction across the full datasets from four published-study cohorts, comprising 769 metagenomic profiles. The heterogeneous inputs included raw paired-end reads, preprocessed host-depleted reads, LC-MS and FTICR-MS matrices, feature annotations and study-specific metadata. Sorghum reanalysis detected 4/4 anchored marker groups and a 2.47-fold Actinobacteria enrichment, alongside treatment-compartment community structure (PERMANOVA R2 = 0.482, p = 0.001). RRMS reanalysis recovered 3/3 anchored features, with beta-diversity separation (p = 0.002) and directionally lower hypotaurine (RRMS-HC log2FC = -1.402; two-sided p = 0.0939). CRC reanalysis recovered 4/5 canonical markers among the top 20 classifier features and achieved AUC = 0.850. Permafrost reanalysis detected 5/5 anchor clade names in the complete taxonomy table and identified 16,141 of 254,848 microbe-metabolite correlations at FDR < 0.05. These results show that one auditable framework can coordinate heterogeneous inputs.

Comparative perturbation experiments addressed robustness separately from biological case interpretation. The upstream pipeline completed reliably, but downstream validity depended on the backend and prompt granularity, with Qwen emerging as the most robust backend overall. DeepSeek and Qwen were broadly concordant across prompt-length comparisons, with only beta diversity in the Simple condition differing (p = 0.0055). Under decoy perturbation, Qwen’s alpha diversity, beta diversity and log2FC did not differ significantly (p = 0.2550, 0.1770 and 0.0712). The main instability was therefore confined to a single prompt–endpoint combination, suggesting that prompt detail is more important for preserving downstream structure than decoy noise once the intended analytical path has been recovered.

Ablation experiments tested component necessity rather than output agreement. Removing the pipeline registry, planning loop or per-job manifest degraded agent performance in characteristically different ways, increasing session duration, token consumption or both. The registry provides a compact machine-readable contract between the upstream and downstream layers. The planning loop supports iterative correction before and during execution. The per-job manifest preserves the state needed to trace and reproduce a run. Their non-redundant ablation profiles indicate that each component protects a distinct aspect of agent performance. Thus, reproducible scientific agency appears to require coordinated control of tool discovery, planning and provenance, rather than relying on prompt design or model scale alone.

Although the benchmarks are anchored in metagenomics, the properties that make MetaClaw auditable are not intrinsically specific to this field. The registry contract, network-isolated downstream container and rerunnable job bundle require only a separation between deterministic preprocessing and configurable interpretation, a structure shared by transcriptomic, proteomic, single-cell and integrative multi-omics pipelines. MetaClaw therefore suggests a general design pattern for reproducible agentic bioinformatics, demonstrated here through metagenomic and microbiome multi-omics reproductions. This separation also supports model-backend evolution: newer LLMs can replace current models without changing registered tool flows, skill contracts or the archival structure. We did not, however, benchmark non-microbiome omics, and generality beyond the tested settings remains to be demonstrated empirically.

Several limitations bound these claims. The benchmarks focused on retrospective reproduction through the read-based taxonomic profiling path; functional profiling, assembly, amplicon and deep-learning pipelines were not benchmarked. Complete end-to-end rerunning currently requires FlowHub access, and we did not test third-party reproduction from archived bundles alone or compare MetaClaw head to head with other agents on the same cohorts. Larger prospective benchmarks with preregistered outputs are needed to test generalization beyond the four evaluated settings. In particular, the evidence for the design pattern’s generality is architectural rather than empirical; extension to transcriptomic, proteomic or clinical multi-omics agents will require dedicated benchmarks. Besides, we are planning to apply MetaClaw as agentic solutions to end-to-end metagenomics and multi-omics analysis in various scenarios, ranging from small research groups to large institute centers.

## 4 Conclusion

MetaClaw couples versioned FlowHub execution to customizable OpenClaw downstream skills through a single YAML pipeline registry. Security and reproducibility are enforced through network-isolated downstream execution and per-job bundles containing FlowHub specifications, skill scripts, pinned Dockerfiles, parameters and outputs. Across the full datasets from four published cohorts, MetaClaw processed 769 metagenomic profiles and recovered 4/4 sorghum marker groups, 3/3 RRMS features, 4/5 canonical CRC markers among the top 20 classifier features and 5/5 permafrost anchor clade names. Across 45 model-by-prompt runs, all backends completed upstream processing, whereas downstream validity depended on model and instruction detail; three decoy-tested endpoints showed no significant differences. Across 48 ablation sessions, each component removal produced a distinct degradation. Relative to registry removal, the intact scaffold reduced time by 32%, tool calls by 29% and token cost by 44%. These findings connect modular architecture, secure rerunnable provenance, conclusion-level reproduction, perturbation robustness and measurable resource efficiency. Current evidence remains limited to retrospective, read-based metagenomic and microbiome multi-omics reproduction; broader claims require prospective benchmarks and independent reruns from archived bundles.

## 5 Methods

### 5.1 System overview

MetaClaw is implemented as a thin LLM-driven gateway positioned between a chat interface and two execution backends—a cloud-side upstream backend and a host-side downstream backend—coupled through a shared job workspace on the host filesystem (**Fig. 1**). The gateway runs as a persistent process on a Linux host and, by design of OpenClaw, is agnostic to the chat platform: the same agent can be reached from WhatsApp, Slack, Telegram or a command-line client without code changes. At start-up, the gateway loads several metagenomics-oriented workspace files that define the agent’s behaviour while separating concerns: SOUL.md (persona and boundaries), AGENTS.md, TOOLS.md (the host commands and container entry points that the agent may call), HEARTBEAT.md (the periodic self-check for long-running jobs), and IDENTITY.md, USER.md and MEMORY.md (identity, user preferences and durable memory). Three task workers use these files at runtime: a Planner Agent maps a user request to a pipeline, a Skill Loader injects the relevant SKILL.md documents into the LLM context, and an Orchestrator executes the resulting plan against the upstream and downstream backends.

Users may reach the gateway through the web deployment at https://www.bioline.cloud or a configured OpenClaw client. Full upstream execution requires access to a FlowHub project; downstream-only analyses can begin from precomputed tables on the host.

The two backends are deliberately heterogeneous. Standardized upstream tasks—including quality control, host-read decontamination, taxonomic profiling and optional assembly—run as versioned FlowHub flows. Downstream statistical analysis, machine learning, visualization and reporting run locally in pinned OpenClaw containers. The backends exchange only staged outputs through a partitioned host workspace; the LLM remains in the gateway and does not execute within either backend.

Note that the paper uses “reproduce” both for scientific reproduction of conclusions and computational reproducibility of a run, and that the four benchmarks test the former while the archival design targets the latter.

### 5.2 Upstream execution on FlowHub

MetaClaw invokes registered FlowHub flows through its command-line client using a plan–submit– poll–finalise lifecycle. Before submission, the user selects a FlowHub project and provides the absolute path to data already present in its filesystem. Planning checks the selected flow, enumerates inputs and previews file bindings for user confirmation; submission then applies any parameter overrides and starts the versioned workflow.

Long-running jobs are polled asynchronously, and each status snapshot is archived. After successful completion, finalisation stages the declared outputs in the host workspace and writes a completion marker before downstream skills can run. Cohort-scale jobs use the same lifecycle in per-sample batch mode: input files are grouped by sample, progress is tracked separately, and finalisation occurs only after all required samples succeed.

### 5.3 Downstream execution in local OpenClaw skills

Downstream analysis is organized as a catalogue of OpenClaw skills. Each skill resides in skills/<NAME>/ and comprises a SKILL.md document, containing YAML-fronted instructions rendered into the LLM context by the Skill Loader, and a reference script under scripts/. The reference script is not executed directly; instead, it serves as a documented starting point that the LLM adapts to the dataset. This reference–customize–archive–execute pattern allows each skill to encode domain best practice while retaining flexibility for cohort-specific properties, such as group balance, covariates and sample size, that are known only after the data have been inspected.

All skills use one of three pinned container images. openclaw/downstream is the default image and contains the Python and R stacks required for statistics, classical machine learning, visualization and reporting; openclaw/downstream-dl adds PyTorch and protein language-model dependencies for deep-learning skills; and openclaw/base is a minimal image for lightweight text-processing skills and adapters to external tools. At job time, the Orchestrator starts a long-running container, mounts the job folder and copies the relevant SKILL.md files and reference scripts. It then traverses the skill catalogue and executes the latest LLM-generated or reference script for each skill. Exit codes, log paths and timestamps are written to the reproducibility folder. The container is destroyed at job completion, so no state persists between runs except that archived in the job folder.

### 5.4 Pipeline registry and the upstream***–***downstream contract

A single YAML file is the only coupling point between cloud-side upstream execution and host-side downstream analysis. Each entry declares the FlowHub flow keyword, pipeline-level default parameters to merge into the FlowHub specification, the downstream container image, the catalogue of downstream skills and a timeout_minutes budget used by the heartbeat to flag stalled jobs. Per-sample batch behavior, when applicable, is declared in the same entry under upstream.batch.mode. Adding a pipeline, changing a FlowHub flow version or substituting a downstream container image therefore requires modification of only the registry. Adjacent layers—the FlowHub flows, Docker images and skills—do not refer to one another directly. Per-job customizations do not overwrite the registry, but are stored in a separate parameter-overrides YAML file.

### 5.5 Reproducibility and security

Reproducibility is implemented as an auditable record rather than a post hoc description. MetaClaw archives the FlowHub workflow version, input identifiers, effective parameters and monitoring records; for downstream analysis, it preserves each invoked SKILL.md file, the executed scripts and the pinned container definitions. A rerun script links these materials into a per-job bundle intended to make the analytical procedure inspectable and reproducible.

The same architecture supports a conservative security model. The LLM resides only in the host gateway and does not enter either backend: the upstream flow is a closed FlowHub container chain receiving a static JSON specification, and the downstream container receives a saved script rather than free-form instructions. This design closes a common prompt-injection path into the analytical runtime. The downstream container is launched with --network none and --cap-drop ALL, preventing analysis scripts from exfiltrating data or escalating privileges. Raw sequencing reads remain on the standalone FlowHub filesystem throughout the job, reducing the host attack surface and supporting cohorts whose data-handling agreements prohibit copying reads from controlled storage. In this sense, security is part of the reproducible-agent paradigm: the boundaries that make a run inspectable also constrain where untrusted content and LLM-generated code can act.

### 5.6 Benchmark design

We selected four previously published shotgun-metagenomic studies as reproduction targets. For each study, raw FASTQ reads were placed on FlowHub, whereas published metadata tables and task-specific auxiliary files were provided separately as benchmark inputs. These auxiliary files included, for example, the metabolite abundance and annotation matrices used in Benchmarks 2 and 4. The agent received a free-text request paraphrasing the original paper’s analytical goal and was not given analysis scripts. The four benchmarks exercised the read-based taxonomic profiling path—quality control, host-read removal and MetaPhlAn 4 profiling. Functional profiling, assembly, amplicon and deep-learning pipelines described in Section 2.1 are framework capabilities and were not evaluated in these reproductions. Unless otherwise stated, the reproductions used the Qwen 3.7-Max model through Alibaba Cloud’s Bailian API, with the qwen3.7-max backend in high-thinking mode.

#### 5.6.1 Benchmark 1 **—** Sorghum rhizosphere drought-response metagenomics

This benchmark was based on the sorghum rhizosphere and bulk-soil metagenomic study of Xu et al. [21], which investigated drought-induced microbiome restructuring and reported enrichment of Actinobacteria and iron-metabolism-associated taxa under drought stress. The complete cohort contained 47 paired-end shotgun metagenomes from BioProject PRJNA657940, sequenced on the Illumina NovaSeq 6000 platform: 11 drought-rhizosphere, 12 watered-rhizosphere, 12 drought-bulk-soil and 12 watered-bulk-soil samples. The reproduction target was to recover drought-associated taxonomic shifts, compartment-level community separation and Actinobacteria/iron-associated marker enrichment.

#### 5.6.2 Benchmark 2 **—** RRMS saliva microbiome**–**metabolome profiling

This benchmark was based on the relapsing-remitting multiple sclerosis saliva microbiome and metabolomics study of Fitzjerrells et al. [22], which reported oral dysbiosis, reduced early colonizers including Streptococcus and decreased hypotaurine levels in RRMS patients. The complete metagenomic cohort contained 99 paired-end saliva shotgun metagenomes from BioProject PRJNA1090491, sequenced on the DNBSEQ-G400 platform (50 healthy controls and 49 RRMS), and LC-MS metabolite abundance data for 20 samples. The reproduction target was to recover HC-versus-RRMS microbiome community differences, Streptococcus-associated taxonomic shifts and the targeted hypotaurine depletion signal.

#### 5.6.3 Benchmark 3 **—** Colorectal cancer stool metagenomic biomarker classification

This benchmark was based on the French cohort from Zeller et al. [23], which demonstrated the potential of fecal metagenomic species profiles for colorectal cancer detection. The complete cohort contained 156 stool shotgun metagenomes from ENA project PRJEB6070: 61 healthy controls, 42 adenoma samples and 53 CRC samples. The benchmark used the published preprocessed reads after adapter screening and human host-read removal. The reproduction target was to recover CRC-associated community structure, detect canonical CRC-associated species and train a microbiome-based classifier for CRC versus non-CRC.

#### 5.6.4 Benchmark 4 **—** Stordalen Mire permafrost thaw-gradient microbiome**–**metabolome analysis

This benchmark was based on the Nature Microbiology study of microbiome-metabolite linkages across Palsa, Bog and Fen habitats in the Stordalen Mire permafrost thaw gradient [24]. The complete taxonomic output contained 467 peat shotgun metagenomic profiles from BioProject PRJNA386568. Habitat-aware analyses used 154 labelled profiles (33 Palsa, 63 Bog and 58 Fen) and incorporated paired FTICR-MS metabolite abundance, annotation and sample-metadata tables for multi-omics integration. The reproduction target was to recover habitat-associated microbial alpha- and beta-diversity patterns, detect the expected greenhouse-gas-associated marker names, test habitat-structured metabolite features and evaluate correlative microbe-metabolite associations.

#### 5.6.5 Evaluation metrics

Agent performance was evaluated at three levels: workflow completion, analytical validity, and concordance with publication-anchored biological targets.

Workflow completion was assessed by determining whether the agent generated the essential intermediate and final outputs required for each benchmark. These included a merged species-level abundance table suitable for downstream analysis, diversity-analysis results, differential-abundance tables, visualizations, and a final analysis report. Benchmark-specific outputs were also required. For the CRC benchmark, these included random-forest classifier metrics and biomarker rankings; for the RRMS benchmark, these included metabolite-level differential analysis and targeted testing of hypotaurine and taurine.

Analytical validity was assessed using task-specific quantitative outputs. Across all benchmarks, alpha diversity was evaluated using standard indices, including Shannon diversity, Simpson diversity, and observed species richness. Beta diversity was evaluated using Bray–Curtis dissimilarity, principal-coordinate analysis (PCoA), and PERMANOVA tests for group-associated community differences. Differential-abundance analyses were evaluated according to the presence, direction, nominal significance, and multiple-testing-adjusted significance of study-relevant taxa. For the RRMS benchmark, metabolite-level results were evaluated using log-transformed abundance comparisons and targeted tests of hypotaurine and taurine. For the CRC benchmark, classifier performance was evaluated using the area under the receiver operating characteristic curve (AUC-ROC), sensitivity, specificity, positive predictive value, and negative predictive value.

Biological concordance was evaluated against predefined publication-anchored targets for each benchmark. For the sorghum drought benchmark, concordance was assessed by recovery of drought-associated *Actinobacteria* enrichment and overlap with expected drought- or iron-associated microbial groups. For the RRMS benchmark, concordance was assessed by recovery of *Streptococcus*-associated oral dysbiosis and reduced hypotaurine levels in RRMS samples. For the CRC benchmark, concordance was assessed by detection of canonical CRC-associated markers before classifier modeling, including *Fusobacterium nucleatum*, *Peptostreptococcus stomatis*, *Parvimonas micra*, *Gemella morbillorum*, and *Porphyromonas asaccharolytica*, together with the predictive performance of the random-forest classifier.

### 5.7 Robustness evaluation across models, prompts and input perturbations

To quantify stability under model, prompt and input perturbations, we performed three complementary robustness evaluations while holding upstream preprocessing fixed. The first used a 3 × 3 factorial design on the sorghum rhizosphere benchmark from Xu et al. [21], with 10% randomly sampled reads for efficiency. Three LLM models—Doubao (doubao-seed-2-0-pro-260215), DeepSeek (deepseek-v4-pro) and Qwen (qwen3.7-max)— were crossed with three prompt specifications (Full, Medium and Simple).

The Full prompt provided the most detailed guidance, including the complete analytical workflow, exact statistical parameters (rarefaction depth, permutation count and FDR threshold), an explicit multifactor PERMANOVA formula, visualization requirements, output naming conventions and quality-control criteria.

The Medium prompt retained the same core analytical goals and named the statistical tests, but omitted parameter values, the multifactor model and detailed output formatting.

The Simple prompt stated only the high-level task and included no explicit statistical, visualization or file-organization instructions.

We ran five independent replicates per condition, yielding 45 runs. The upstream stage was identical in all runs and consisted of read quality control, host-read removal and MetaPhlAn 4 taxonomic profiling. Panel A reports valid completion across pipeline stages; a run was considered valid only when the expected output existed and passed stage-specific content checks, including production of sample-level rather than taxon-level alpha- and beta-diversity outputs. Panel B compares DeepSeek and Qwen using pooled raw replicate-level values across shared valid samples and features, with two-sided Mann–Whitney U tests and exclusion of zero-Shannon alpha-diversity rows before pooling. In the second evaluation, Qwen was used to compare Full, Medium and Simple prompts. For alpha and beta diversity, values were pooled across sample labels shared by all prompt groups; for differential abundance, values were pooled across species or features shared by all groups. Zero-Shannon rows and unclassified-only features were excluded, and differential-abundance analysis was restricted to species-level features. Pairwise prompt contrasts used two-sided Mann–Whitney U tests with Holm correction within each metric. In the third evaluation, Qwen was used for the decoy test. Raw replicate-level values were pooled across shared valid sample labels for alpha and beta diversity and across shared species or features for log2FC. Unclassified-only features were excluded, and differential abundance was restricted to species-level features. Each run was checklist-valid only when the expected output existed and passed stage-specific content checks.

### 5.8 Ablation of MetaClaw’s scaffold components

To estimate how much of MetaClaw’s performance derives from its scaffolding rather than the underlying language model, we created three ablated variants of the agent workspace and applied them to the first benchmark (sorghum drought, using 10% reads from 20 random samples for efficiency). Each variant removed one scaffold component while leaving the others intact.

“-Plan” stripped the deliberative planning prompts in AGENTS.md and replaced them with immediate-action stubs, so the agent picked a pipeline and ran it without inspecting the data first or confirming with the user.

“-Registry” reduced “pipelines.yaml” to a six-line skeleton, forcing the agent to derive flow keywords, port bindings, batch detection and default parameters from inspection of the FlowHub file system.

“-Manifest” suppressed every write to “stage/manifest.json” and the per-skill exit-code log, so downstream skills could no longer rely on the structured hand-off index that their SKILL.md files still pointed them at.

An untouched workspace served as the baseline arm. Each arm was run 12 times on the same 20-sample Xu et al. [21] subset using the same Qwen 3-Max backend, yielding 48 independent sessions.

Output completeness for each session was scored against a content-validated checklist of 15 deliverables grouped into five weighted categories: the merged taxonomic table; diversity outputs (alpha table, alpha boxplot and Bray–Curtis PCoA plot); differential abundance outputs (rhizosphere and bulk-soil tables and volcano plot); figures (top-30 heatmap and phylum bar plot); and the narrative report. A match required both a filename check and a content check: a MetaPhlAn-style taxonomic column for the species table, numeric columns for the diversity table, paired log2FC and p-value columns for differential-abundance tables, a valid PDF or PNG header for figures and a heading-rich body for the Markdown report. Per-category scores were combined as a weighted average to give session-level completeness on a scale from 0 to 1.

Wall-clock duration, tool invocations and total token cost were read directly from the per-session ledger, which records new-input, output, cache-created and cache-hit tokens. For each metric, the 12 replicates per arm were sorted and the smallest and largest values were removed, leaving 10 values per arm for descriptive statistics and significance testing. Trimming the two extremes reduces the influence of the two most extreme replicates while retaining the central distribution. Results are reported as the mean ± standard error of the mean (SEM). Each ablation arm was compared with baseline using a two-sided Mann–Whitney U test on the trimmed n = 10 samples. No multiple-comparison adjustment was applied because the three contrasts were pre-specified.

### 5.9 Use of large language models in manuscript preparation

Claude and OpenAI Codex were used to assist with grammar and readability of the manuscript text. All scientific content, data analysis, and interpretation are the authors’ own work, and the authors take full responsibility for the accuracy and integrity of the manuscript. This is distinct from MetaClaw’s use of LLM backends (Qwen3.7-Max, DeepSeek-V4-Pro, Doubao-Seed-2.0-Pro), which is the technology under study rather than a tool used in preparing this text.

## Acknowledgements

The authors thank Fei Wang, Hang Wang, Feng Zhang and Yang Zhao for their support with tools and flows on FlowHub.

## Funding

This work was partially supported by the National Key R&D Program of China (Grant Nos. 2023YFA1800900 and 2018YFC0910502) and the National Natural Science Foundation of China (Grant Nos. 32071465, 31871334 and 81827901).

## Author contributions statement

**H.Z.** and **Z.L.** contributed equally to this work. **H.Z.**: Conceptualization, Methodology, Validation, Formal analysis, Investigation, Writing - Original Draft, Visualization. **Z.L.**: Conceptualization, Methodology, Software, Formal analysis, Investigation, Writing - Original Draft, Visualization. **P.N.P.L.**: Software, Formal analysis, Investigation, Writing - Original Draft, Visualization. **Z.W.**: Validation, Formal analysis, Investigation, Writing - Review & Editing. **L.Z.**: Validation, Writing - Review & Editing. **W.L.**: Software, Writing - Review & Editing. **P.D.**: Software, Writing - Review & Editing. **X.J.**: Conceptualization, Resources, Writing - Review & Editing, Supervision, Project administration. **K.N.**: Conceptualization, Resources, Writing - Review & Editing, Visualization, Supervision, Project administration, Funding acquisition.

## Competing interests

The authors declare no competing interests.

## Data availability

The raw shotgun metagenomic sequencing reads reanalyzed in this study are publicly available: sorghum rhizosphere reads under BioProject PRJNA657940; RRMS saliva reads under BioProject PRJNA1090491; colorectal cancer stool reads under ENA project PRJEB6070; and permafrost thaw-gradient reads under BioProject PRJNA386568. Matched metabolomics data (LC–MS for the RRMS benchmark; FTICR-MS for the permafrost benchmark) and sample metadata were retrieved from the EMERGE Database repository reported in the original publication: https://emerge-db.asc.ohio-state.edu/datasources/0141_Wilson-etal-2022-STOTEN_ICR-plants. Processed data generated in this study, including merged taxonomic abundance tables, diversity and differential-abundance results, robustness- and ablation-analysis outputs, are available in the Supplementary Information. Source data are provided with this paper.

## Code availability

Source code for MetaClaw, including the OpenClaw agent workspace files (SOUL.md, AGENTS.md, TOOLS.md, HEARTBEAT.md, IDENTITY.md, USER.md, MEMORY.md), the pipeline registry (pipelines.yaml), the SKILL.md contracts and reference scripts for each downstream skill, the Dockerfiles for the three pinned container images (openclaw/downstream, openclaw/downstream-dl, openclaw/base), and the benchmark and ablation configuration files, is available at https://github.com/BGI-AI-models/MetaClaw/ under the MIT License. An online deployment of MetaClaw is available at https://www.bioline.cloud, allowing users to access the framework through a web interface. The upstream stage additionally depends on the FlowHub platform; full end-to-end execution of the pipelines requires FlowHub access, as noted in the Discussion. The archived FlowHub specifications, tool contracts and downstream bundle remain inspectable without FlowHub access.

## Notes

### Competing Interest Statement

The authors have declared no competing interest.

### Summary of Updates

This revision expands and clarifies the validation of MetaClaw. We extended the benchmark to the full cohorts of four published studies, comprising 769 metagenomic profiles, and updated the associated analyses, figures, and quantitative results. MetaClaw recovered 4/4 sorghum marker groups, 3/3 relapsing-remitting multiple sclerosis features, 4/5 canonical colorectal cancer markers, and 5/5 permafrost marker groups. We also redrew the Study 4 sample-level abundance boxplots, applied consistent statistical testing for alpha diversity, and annotated the overlap analyses with the identities and recovery status of study-specific markers. The robustness and ablation evaluations were updated to include 45 model-by-prompt runs and 48 ablation sessions. In addition, we revised the Abstract, Introduction, and Methods to clarify the study rationale and technical contribution; clarified the high-level use of FlowHub; discussed the limitations of general-purpose workflow engines such as Snakemake; added citations and analytical motivations for the four case studies; incorporated the online platform at www.bioline.cloud; removed unnecessary cohort-specific data descriptions; and updated the supporting references. Together, these revisions provide additional validation and methodological detail while leaving the central conclusions of the study unchanged.

